# Maternal care of heterozygous Dopamine Receptor D4 knockout mice: Differential susceptibility to early-life rearing conditions

**DOI:** 10.1101/761049

**Authors:** Jelle Knop, Marinus H. van IJzendoorn, Marian J. Bakermans-Kranenburg, Marian Joëls, Rixt van der Veen

**Affiliations:** Dept. Translational Neuroscience, Brain Center Rudolf Magnus, University Medical Center Utrecht, Utrecht University, Utrecht, The Netherlands; Faculty of Social and Behavioural Sciences, Leiden University, Leiden, The Netherlands; Dept. of Psychology, Education and Child Studies, Erasmus University Rotterdam, Rotterdam, The Netherlands; Primary Care Unit, School of Clinical Medicine, University of Cambridge, Cambridge, United Kingdom; Clinical Child & Family Studies, Vrije Universiteit Amsterdam, Amsterdam, Netherlands; University of Groningen, University Medical Center Groningen, Groningen, The Netherlands

## Abstract

The differential susceptibility hypothesis proposes that individuals who are more susceptible to the negative effects of adverse rearing conditions may also benefit more from enriched environments. Evidence derived from human experiments suggests the lower efficacy dopamine receptor D4 (*DRD4*) 7-repeat as a main factor in exhibiting these for better and for worse characteristics. However, human studies lack the genetic and environmental control offered by animal experiments, complicating assessment of causal relations. To study differential susceptibility in an animal model, we exposed *Drd4*^*+/-*^ mice and control litter mates to a limited nesting/bedding (LN), standard nesting (SN) or communal nesting (CN) rearing environment from postnatal day (P) 2-14. Puberty onset was examined from P24-P36 and adult females were assessed on maternal care towards their own offspring. In both males and females, LN reared mice showed a delay in puberty onset that was partly mediated by a reduction in body weight at weaning, irrespective of *Drd4* genotype. During adulthood, LN reared females exhibited characteristics of poor maternal care, whereas dams reared in CN environments showed lower rates of unpredictability towards their own offspring. Differential susceptibility was observed only for licking/grooming levels of female offspring towards their litter; LN reared *Drd4*^*+/-*^ mice exhibited the lowest and CN reared *Drd4*^*+/-*^ mice the highest levels of licking/grooming. These results indicate that both genetic and early-environmental factors play an important role in shaping maternal care of the offspring for better and for worse.

## Introduction

### 1.1. Differential susceptibility

Parental care is essential for survival and development of newborn mammals, including humans. Variations in parental care substantially contribute to the environmental variability experienced by offspring. Extensive evidence indicates that poor parental care can contribute to increased vulnerability to develop later-life psychopathology in humans and impaired cognitive performance in rodents (Gunnar et al., 2015; Krugers and Joëls, 2014). This vulnerability crucially depends on a complex cross-talk between an individual’s genetic makeup and rearing environment (Nugent et al., 2011). While the genetic background of some individuals is related to a vulnerable phenotype in the face of early-life adversity, others appear to be more resilient. Interestingly, individuals who are genetically more susceptible to the detrimental consequences of negative (rearing) conditions may also experience greater benefits from a positive and stimulating (rearing) environment (Belsky and Van IJzendoorn, 2017; Ellis et al., 2011). This crossover effect *for better and for worse*, also called differential susceptibility, is supported by studies investigating the role of human allelic variation across a variety of susceptibility genes (Bakermans-Kranenburg and Van IJzendoorn, 2015).

An example of such differentially susceptibility concerns the exon III 7-repeat polymorphism of the D2-like dopamine receptor D4 gene (*DRD4-7R*). In humans, this variant has been associated with reduced gene expression and efficiency (Asghari et al., 1995; Schoots and Van Tol, 2003) and acts as a susceptibility marker of dopamine-related genes (Bakermans-Kranenburg and Van IJzendoorn, 2015). Carriers of this variant have an increased risk of developing externalizing problems in relation to parental insensitivity (Bakermans-Kranenburg and van IJzendoorn, 2006) and chronic stress (Zandstra et al., 2018). However, these individuals also benefitted most from enhanced positive parenting (Bakermans-Kranenburg et al., 2008b). Meta-analytic evidence further supports an important role of dopamine-related genes in moderating susceptibility to both positive and negative rearing environments (Bakermans-Kranenburg and van IJzendoorn, 2011). Of note, the *DRD4* also plays a role in moderating parental care itself (Leerkes et al., 2017; van IJzendoorn et al., 2008).

### 1.2 Rodent models of impoverished or enriched rearing environments

Studying differential susceptibility in humans is hampered by random genetic variability. Moreover, it is often difficult to randomly allocate individuals to specific environments while also taking genotype into account. Therefore, we set out to study the causal contribution of decreased *Drd4* functioning to differential susceptibility with a truly randomized experiment in rodents, allowing strict control for both genetic variation and environmental factors (Knop et al., 2017). By using *Drd4*^*+/-*^ mice, we aimed to mimic the reduced *DRD4* efficiency observed in human *DRD4-7R* allele carriers.

We selected two rodent models developed to chronically induce alterations in the quality and quantity of parental care received by offspring. First, limited availability of nesting and bedding (LN) material to a mouse dam was used to induce an adverse early life environment; this model increases unpredictability of maternal care received by the pups (Davis et al., 2017; Knop et al., 2019; Molet et al., 2016), leading to increased corticosterone levels in pups (Rice et al., 2008) and altered offspring development and behavior in adulthood (Bonapersona et al., 2019; Walker et al., 2017). Second, as beneficial and stimulating social rearing environment we selected a communal nesting (CN) condition, where two or more dams share care-giving behavior towards multiple litters (Branchi et al., 2006). In this condition, pups experience higher levels of nest occupancy by at least one dam (Branchi et al., 2013; Knop et al., 2019) and can interact with peers as well as siblings. Mice reared in communal nesting conditions exhibit various neurobiological and behavioral characteristics that are indicative of improved social competences (Branchi and Cirulli, 2014).

### 1.3 Outcome parameters

In line with a previous study (Knop et al., 2019), we focused on timing of puberty onset, a key moment in development that is malleable by environmental influences as part of an adaptive reproductive strategy (Belsky et al., 1991). Although adverse rearing conditions in females are linked to accelerated pubertal onset in humans (Belsky et al., 2015) and rats (Cowan and Richardson, 2018), such effects have not yet been observed in mice (Knop et al., 2019). In human males, adverse rearing conditions had no effect on puberty onset (Grassi-Oliveira et al., 2016), while puberty onset in male rodents was either unaffected or delayed (Biagini and Pich, 2002; Cowan and Richardson, 2018; Knop et al., 2019). However, rodent models of early-life adversity (ELA) invariably decrease body weight gain, which is an important mediator of puberty onset. Therefore, it is unclear whether the delayed puberty onset observed in ELA reared animals is the result of decreased body weight gain or whether a *relative* acceleration irrespective of body weight exists in rodents as well.

A second outcome was maternal care provided by female offspring. Preclinical studies allow for feasible, controlled intergenerational studies on maternal care and extensive evidence suggests that alterations in maternal care may be transmitted across generations (Meaney, 2001). Variations in levels of licking/grooming (LG) behavior and arched-back nursing (ABN), core features of positive parenting in rodents, have been shown to affect corticosterone reactivity, hippocampal development and maternal care of the offspring (Meaney, 2001). In addition, the limited bedding/nesting model, which evokes changes in maternal care, results in aberrant patterns of maternal care of the offspring (Roth et al., 2009), whereas mice reared in a communal nesting condition display improved maternal behavior towards their own pups (Curley et al., 2009). Taken together, these studies highlight the importance of maternal care for offspring development, as well as the potential of maternal care to be shaped by the early-life environment, contributing to the intergenerational transmission of social behavior.

In this study, we tested heterozygous *Drd4* knock-out (*Drd4*^*+/-*^) mice and control litter mates on susceptibility to both adverse (LN) and enriched (CN) rearing environments to model differential susceptibility in mice. Animals were examined on i) puberty onset, to track early development, ii) maternal care, as an indicator of transgenerational effects and iii) basal corticosterone levels, to investigate involvement of the hypothalamic-pituitary-adrenal-axis (HPA-axis) in differential susceptibility. Although puberty onset would be hypothesized to be accelerated in LN and delayed in CN reared animals according to life history theory, previous findings indicate that the opposite may be true in mice due to the strong effects of body weight. LN reared mice were hypothesized to display poor maternal care, whereas CN reared mice were hypothesized to show enhanced maternal care. To confirm differential susceptibility, these effects would have to be amplified in, or exclusive to, *Drd4*^*+/-*^ mice.

## 1. Materials & Methods

### 2.1 Animals & Housing

B6.129P2-*Drd4*^*tm1Dkg*^/J (*Drd4*^*+/-*^) mice (Rubinstein et al., 1997) were originally obtained from the Jackson Laboratory (Bar Harbor, Maine, USA) and bred in-house with C57BL/6JOlaHsd (breeding colony, originally obtained from Harlan, France) mice for at least 4 generations before experiments started. All breeding was performed in our own animal facility. Wild-type (wt) female C57BL/6 mice were allowed to breed with male *Drd4*^*+/-*^ mice to generate *Drd4*^*+/-*^ F1 offspring and *Drd4*^+/+^ control litter mates. *Drd4*^*+/-*^ mice are viable, healthy and visually indistinguishable from control animals. Between postnatal day 2 and 14 (P2-14), dam and litter were exposed to a limited nesting/bedding (LN), standard (SN) or communal nesting (CN) condition. A total of 129 female and 116 male F1 offspring obtained from 40 breedings was used to assess puberty onset and, in females (n = 75), maternal care of this generation (see Fig 1. for a timeline of the experiment). Puberty onset and F1 maternal care were scored by a trained experimenter blind to rearing condition and genotype of the animals. Mice had *ad libitum* access to water and chow and were housed on a reversed LD cycle (lights off 08:00 h, temperature 21-22 °C, humidity 40-60 %). All experiments were performed in accordance with the EC council directive (86/609/EEC) and approved by the Central Authority for Scientific Procedures on Animals in the Netherlands (CCD approval AVD115002016644).

**Figure 1.**
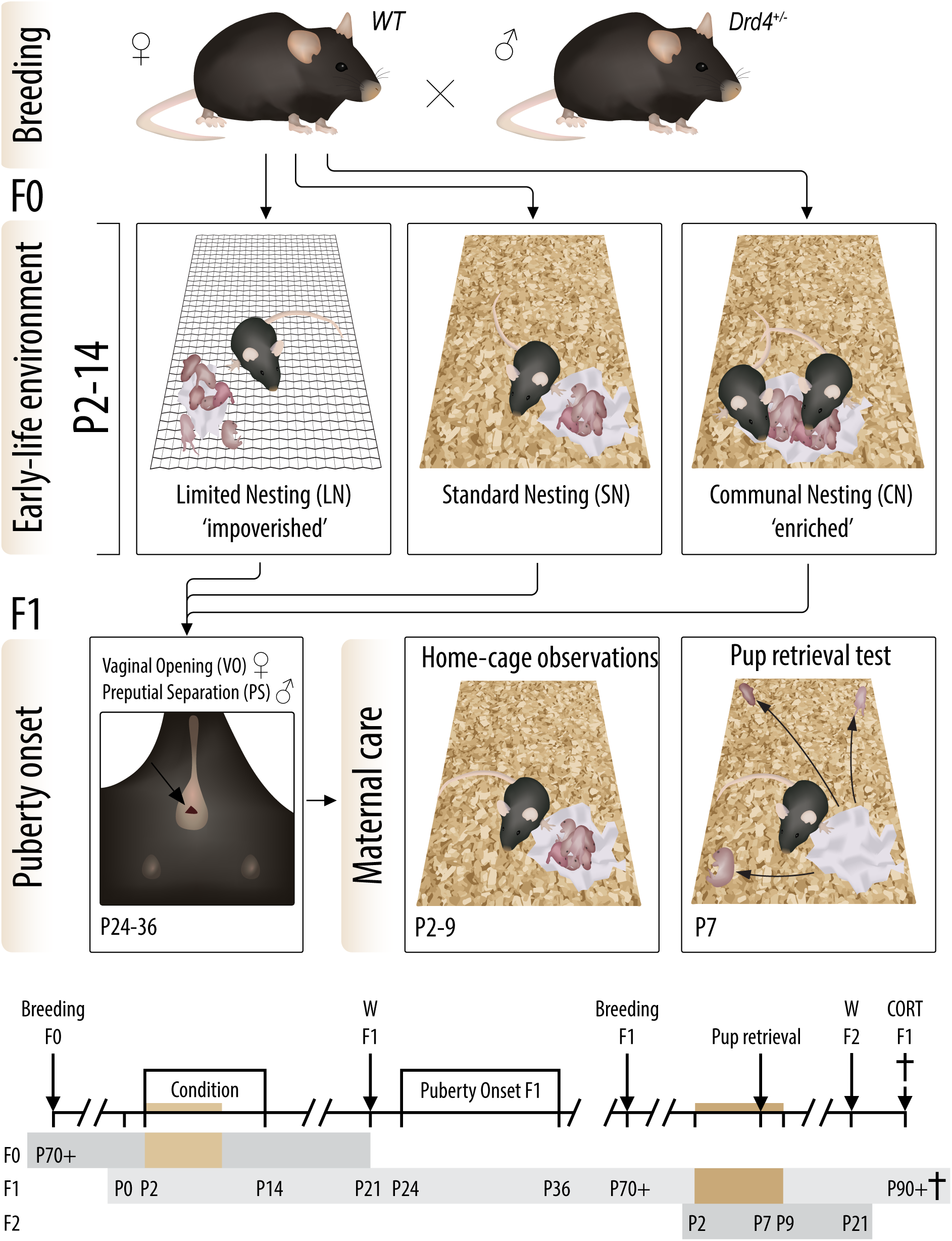
Outline of the experiments. Study design and timeline of the experiment. A wild-type female was paired with a *DRD4*^+/-^ male to obtain litters of mixed genetic background. Experimental time points for each generation of mice are depicted. W = weaning. P = postnatal day. Colored bars indicate periods of home cage maternal care observations.

### 2.2 Breeding conditions

Breeding was performed as described earlier (Knop et al., 2019). In short, one male was paired with two females for 4 days, after which females were co-housed until approximately one week prior to birth. Pregnant dams were then housed in a type II short Macrolon cage (21.5 × 16 cm) with filter top and a Nestlet (5 × 5 cm, Technilab-BMI, Someren, The Netherlands) as nesting material. Daily inspection for the birth of litters was conducted at 09:00 h, assigning the day prior as P0. At P2, dam and litters were weighed and randomly assigned to the LN, SN or CN condition. All litters were culled (or cross-fostered if necessary) to 6-7 pups per litter, with a maximum addition of 1 pup per litter and a minimum of 2 pups of each sex in each litter.

The LN condition consisted of placing the dam and litter in a cage with limited bedding material, covered by a stainless steel wired mesh. In addition only half the regular amount of nesting material was available. In the SN condition, standard amounts of bedding and nesting material were available to the dam. The CN paradigm consisted of co-housing the experimental wt dam (and her genetically heterogeneous F1 litter) with another wt ear-punched dam (and wt litter) in a type II regular Macrolon cage (32×16 cm). The pups of this second mother were marked with surgical marker at P2 and P7 (ArcRoyal, Ireland) to ensure correct allocation of the pups to their mother at the end of communal housing at P14. At P9 and P14, all dams and litters were weighed and provided with clean cages, adding a bit of used bedding material to maintain odor cues. From P14 until weaning at P21, animals were housed in standard nesting conditions. All mice were weighed at weaning and ear punches were obtained to facilitate individual recognition and genotype offspring.

### 2.3 Maternal care observations F0

An instantaneous sampling method (Knop et al., 2019) was used to score maternal behavior of the dams in different conditions. Three 75-minute scoring sessions were performed daily from P2-9 between 06:00-07:30 am (end light phase), 12:00-14:00 pm (mid dark phase) and 16:30-18:30 pm (end dark phase). Red light conditions were used to score during the dark phase sessions. Maternal behavior of each dam was scored every three minutes, leading to 25 observations per session and 75 observations per day for each dam. Maternal behaviors were classified as: arched-back nursing (ABN), passive nursing, licking/grooming pups (LG), nest building, self-grooming on nest, feeding and self-grooming off nest. For observations during which the behavior did not qualify for one of these categories, only on or off nest location of the dam was scored. A Samsung Galaxy Note 4 with Pocket Observer 3.3 software (Noldus, the Netherlands) was used for behavioral scoring, and data was analyzed using Observer XT 10.5 (Noldus, the Netherlands). Both dams in the communal nesting condition were scored separately, using average scores of each pair of dams as an indication of maternal behavior received by the litter.

Assessment of maternal care was performed by looking at i) frequencies of the various maternal behaviors, ii) unpredictability of maternal care and iii) fragmentation, using on/off nest transitions. First, percentage of time spent on the various maternal behaviors was calculated per day (pooling the 3 daily sessions) or circadian phase (pooling over 6 postnatal days) to assess the development over days and circadian rhythmicity of maternal care, respectively. Second, overall unpredictability of maternal behavior was evaluated using the entropy rate of transitions between different maternal behaviors (Molet et al., 2016). The entropy rate is obtained by calculating the probabilities of certain maternal behaviors predicting specific subsequent behaviors, in which higher entropy rates are indicative of higher unpredictability. In addition, unpredictability of maternal care specifically on the nest site was calculated by pooling all off-nest behaviors to enhance representation of the unpredictability rate as experienced by the offspring. Third, the average number of transitions from and to the nest site per observation was used as an index of fragmentation of maternal care (Rice et al., 2008).

### 2.4 Puberty onset F1

As an external measure of puberty onset in males, mice were restrained and gently examined daily form P27-P33 on the potential to fully retract the glans penis and expose the glans penis which was designated as puberty onset (Korenbrot et al., 1977). Female mice were scored daily from P24-P36 for vaginal opening, here taken as sign of puberty onset (Caligioni, 2010). All mice were weighed at puberty onset.

### 2.5 Maternal care F1

During adulthood (>P70), female F1 mice were allowed to breed with a wild-type male as described for F0. All F2 litters were culled/cross-fostered to 6 pups and reared in standard nesting conditions. At P2, P9, P14 and P21, clean cages were provided and animals were weighed. Maternal care observations were performed as described for F0 maternal behavior. At P7 between 10:00-12:00 am, pup retrieval behavior was measured using a 5 minute pup retrieval test as described earlier (Knop et al., 2019). If a dam did not retrieve all three pups within 5 minutes, a latency of 300 seconds was assigned.

### 2.6 Plasma corticosterone levels F1

To measure plasma corticosterone levels, all F1 dams were decapitated in random order between 13:00 – 17:00 at least 3 weeks after weaning of F2 litters. Trunk blood was collected in heparin containing tubes (Sarstedt, The Netherlands) on ice and centrifuged for 10 minutes (13000 rpm) at 4 °C. Plasma was collected and stored at −20 °C until radioimmunoassay (MP Biomedicals, The Netherlands; sensitivity 3 ng/ml).

### 2.6 Statistical analysis

Data are expressed as mean ± SEM. Values deviating >3.29 SD from the mean were defined as outlying and winsorized accordingly (Tabachnick and Fidell, 2007). The entropy rate of one F0 LN dam was winsorized. Data was analyzed using SPSS 23 (IBM) and litter effects in F1 were accounted for using the SPSS complex samples module. However, no effect sizes are provided in this model. In other analyses, eta squared effect sizes (⍰^2^), representing the explained variance relative to the total model variance, are reported. Overall ANOVA statistics are presented in the text, Tukey HSD (main effects) or Sidak (interactions) corrected post-hoc comparisons are depicted in figures.

Greenhouse-Geisser corrected repeated measures ANOVAs with breeding condition as the between-subject factor and postnatal day or observation as within-subject factors were used to analyze F0 maternal behaviors. Maternal behaviors from two observation sessions at P2 were analyzed separately to dissociate acute effects of novel environment exposure from more chronic alterations in maternal care. P2 maternal behavior, entropy rates and fragmentation were analyzed using a one-way ANOVA with breeding condition as the between-subjects factor. Pup retrieval latencies of F1 dams were analyzed using cox regression, as this method is preferred if a subset of animals fails to complete a certain task (Jahn-Eimermacher et al., 2011). All other F1 measures were analyzed using a two-way ANOVA including rearing condition and genotype as independent variables. Pearson correlations were used for correlational data. Mediation analysis was conducted using the PROCESS v3 SPSS macro (Hayes and Preacher, 2014), with rearing condition as a multicategorical independent variable and the SN group as the reference category. The day of puberty onset was used as dependent variable and body weight at weaning and received entropy rates as potential mediators. Significant mediation was assigned when 95% confidence intervals of mediation did not include zero.

## 2. Results

### 3.1 Maternal care F0

The maternal care of mouse dams was affected by environmental condition (Fig. 2). Nesting condition altered arched-back nursing levels (F(2, 36) = 5.69, p = .007, ⍰^2^ = 0.24, Fig. 2a), with increased ABN levels in LN dams compared to individual CN dams. Moreover, nesting condition altered passive nursing levels (F(2, 36) = 5.10, p = .011, ⍰^2^ = 0.22, Fig. 2b). Post-hoc analysis revealed that CN dams spent less time passively nursing their pups compared to SN dams. Together, nesting conditions affected the total nursing levels (i.e., the sum of ABN and passive nursing) displayed by individual dams (F(2, 36) = 14.27, p < .001, ⍰^2^ = 0.44, Fig. 2c). Individual dams in the communal nesting condition exhibited decreased nursing levels compared to both SN and LN dams. In addition, feeding behavior of dams was affected by condition (F(2, 36) = 12.36, p < .001, ⍰^2^ = 0.41, Fig. S1a), with an increase in CN dams compared to both SN and LN animals.

**Figure 2.**
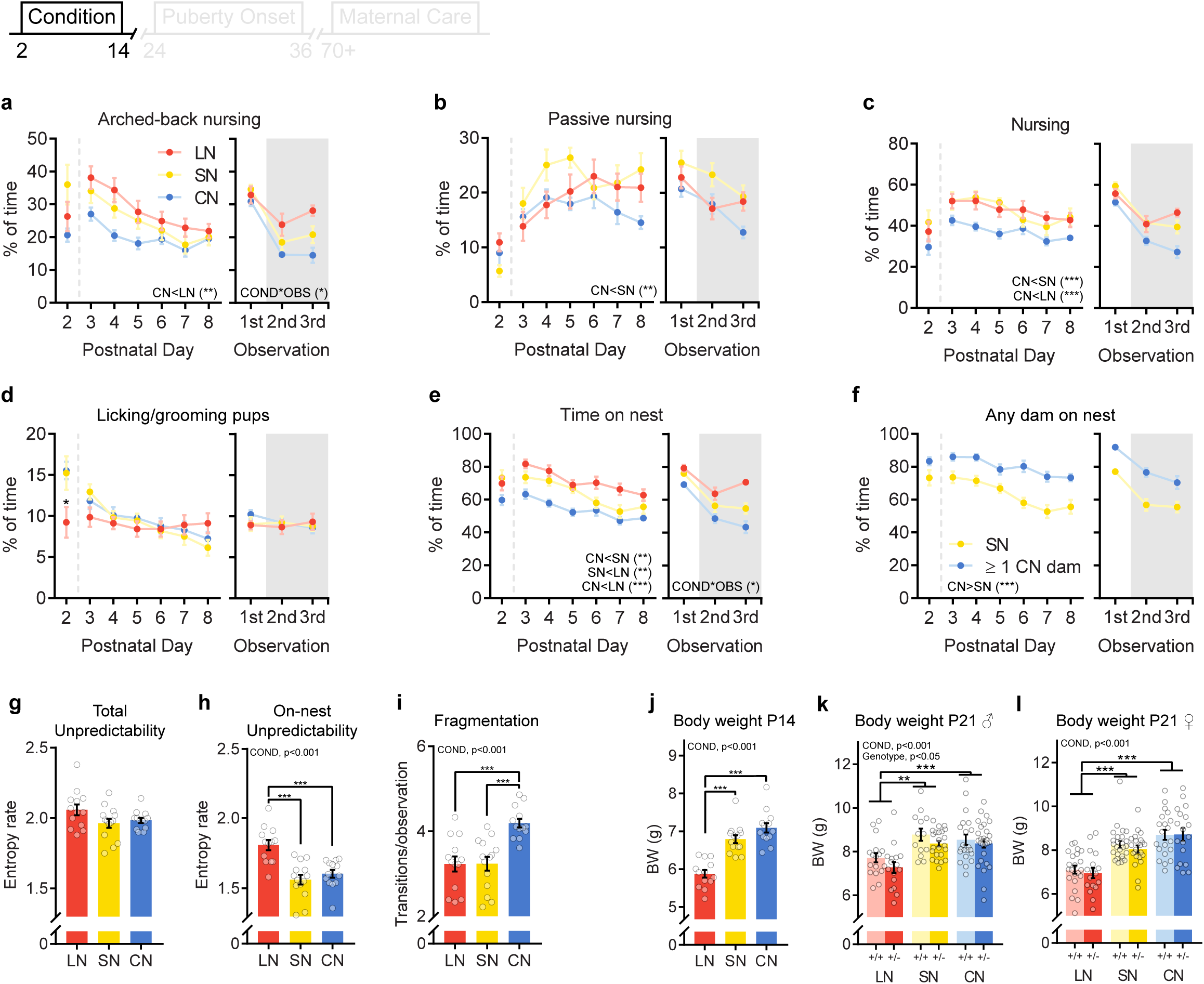
Effect of different housing conditions on F0 maternal care and F1 body weight. (**a**) Arched-back nursing, (**b**) passive nursing, (**c**) total nursing, (**d**) licking/grooming and (**e,f**) time on nest for limited nesting (red, n = 13), standard nesting (yellow, n = 14) and communal nesting (blue, n = 13) dams, depicted over postnatal days (left) and time of the day (right). The shaded area indicates the dark phase of the LD cycle. Data in **f** represents the time on nest by at least one dam from the litters perspective. (**g**) Unpredictability of all scored maternal behaviors and (**h**) unpredictability of maternal care when all off-nest behaviors were combined into one measure. (**i**) Fragmentation (on/off nest transitions) of maternal behavior. Each dot represents one dam and the average of two dams in the CN condition. (**j**) Offspring body weight averaged per litter at postnatal day 14. (**k**) Offspring body weight per individual at weaning for males and (**l**) females. +/+: control, +/-: heterozygous *Drd4*. Group size: ♂: LN +/+: n = 17, LN +/-: n = 16, SN +/+: n = 13, SN +/-: n = 23, CN +/+: n = 22, CN +/-: n = 27; ♀: LN +/+: n = 22, LN +/-: n = 17, SN +/+: n = 26, SN +/-: n = 22, CN +/+: n = 20, CN +/-: n = 18. Asterisks indicate interactions or post-hoc comparisons. \**p* < 0.05, \*\**p* < 0.01, \*\*\**p* < 0.001.

Although condition did not affect licking/grooming behavior from P3-8 (F(2, 36) = 0.13, p = .880, ⍰^2^ = 0.01, Fig. 2d), LG levels were affected more acutely at P2 (F(2, 36) = 4.32, p = .021, ⍰^2^ = 0.19). Post-hoc testing indicated that specifically pups in a LN setting were deprived from LG on this first day of novel environment exposure. The time spent on the nest site differed across conditions (F(2, 36) = 25.95, p < .001, ⍰^2^ = 0.59, Fig 2e); LN dams spent more time on the nest compared to SN and CN dams. This was mostly due to an increase in the time LN dams were engaging in non pup directed behaviors on the nest site (self-grooming and other behavior, see Fig S1). Nest occupancy of individual CN dams was decreased compared to both LN and SN dams. Nevertheless, the nest site in the CN setting had higher levels of nest occupancy by at least one dam compared to the SN condition (F(1, 25) = 76.00, p < .001, ⍰^2^ = 0.75, Fig. 2f). Moreover, circadian rhythmicity of nest occupancy was altered by exposure to different conditions (observation*condition interaction: F(3.51, 64.86) = 2.72, p = .044, ⍰^2^ = 0.06, Fig. 2e, right panel). In particular dams in the LN condition exhibited higher levels towards the end of the dark phase compared to dams in the CN and SN condition. The same pattern was observed in arched-back nursing (observation*condition interaction: F(3.69, 68.31) = 3.40, p = .016, ⍰^2^ = 0.07, Fig. 2e, right panel)

The overall unpredictability of behavior displayed by the dams was not significantly affected by condition (F(2, 37) = 2.70, p = .081, ⍰^2^ = 0.13, Fig. 2g). However, the unpredictability of behavior on the nest site (on nest entropy rates) differed (F(2, 37) = 16.02, p < .001, ⍰^2^ = 0.46, Fig. 2h). Post-hoc comparisons revealed that the LN dams displayed increased unpredictability of maternal care compared to the SN and CN dams. Nesting condition also affected fragmentation of maternal care, measured by the number of transitions from and to the nest site (F(2, 37) = 13.08, p < .001, ⍰^2^ = 0.41, Fig. 2i); individual CN dams exhibited increased fragmentation compared to SN and LN dams.

### 3.2 Body weight F1

While body weight of litters before exposure to different rearing condition was comparable across groups, condition affected litter weight after condition at P14 (F(2, 37) = 32.44, p < .001, ⍰^2^ = 0.64, Fig. 2j). LN reared litters weighed less than both SN and CN litters. At weaning, LN reared mice remained lighter than SN and CN animals, which was found in both males (F(2, 36) = 8.77, p = .003, Fig. 2k) and females (F(2, 37) = 20.60, p < .001, Fig. 2l). In females, heterozygous knock-out of the dopamine receptor D4 did not affect body weight (F(1, 38) = 0.45, p = .505). In contrast, a reduction of body weight was observed in *Drd4*^*+/-*^ males (F(1, 38) = 4.97, p = .032). However, *Drd4* genotype did not interact with rearing condition in either males (F(2, 36) = 0.39, p = .686) or females (F(2, 37) = 0.15, p = .829).

### 3.3 Puberty onset

#### Males

In males, rearing condition affected timing of puberty onset, measured by preputial separation (F(2, 36) = 10.33, p = .001, Fig. 3a). LN reared mice had a delayed puberty onset compared to SN and CN reared mice, but rearing conditions did not interact with *Drd4* genotype (F(2, 36) = 1.84, p = .121). No main effect of *Drd4* genotype on puberty onset was observed (F(1, 37) = 2.83, p = 0.101). Body weight at the day of puberty onset (Fig. 3b) was unaltered by condition (F(2, 34) = 2.20, p = .382) or genotype (F(1, 35) = 0.38, p = .544). Puberty onset negatively correlated with body weight at weaning (r = −0.61, p < .001, Fig. 3c). In males no correlation between received entropy rates during development and puberty onset was found (r = 0.10, p = .278, Fig. 3d). Mediation analysis revealed that in males, the delayed puberty onset found in LN reared mice was partly mediated by the reduced body weight at weaning (95%CI = [0.36, 1.17], Fig. 3e).

**Figure 3.**
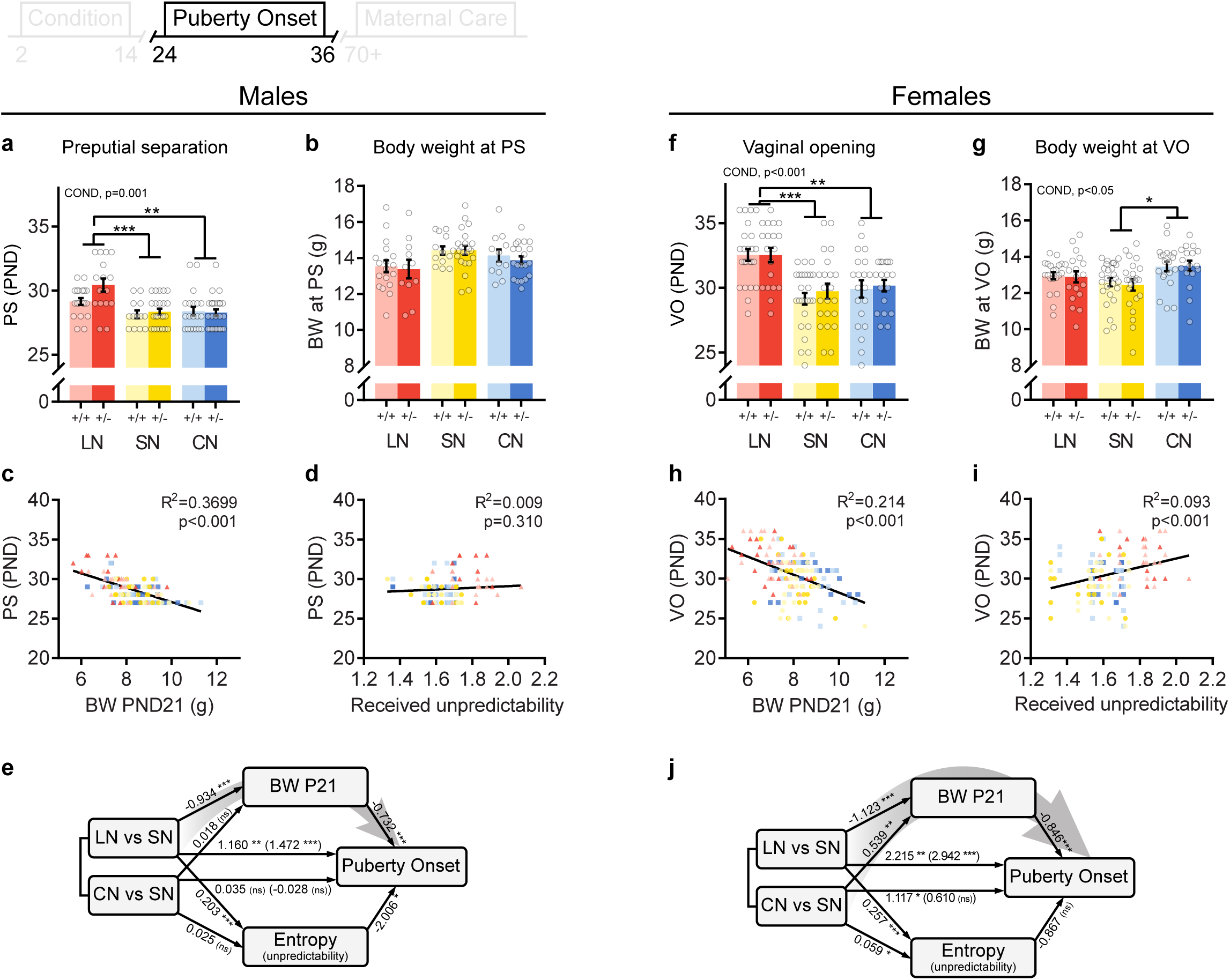
Effects of different rearing conditions on sexual maturation in male and female offspring. (**a,f**) Puberty onset in male (preputial separation) and female (vaginal opening) mice. (**b,g**) Body weight at puberty onset. (**c,h**) Body weight at weaning negatively correlated with puberty onset in both males and females, whereas (**d,i**) received on-nest unpredictability rates during rearing positively correlated with puberty onset only in females. (**e,j**) Graphical representation of mediation models. Numbers represent estimated model coefficients, direct effects are depicted in parenthesis. Grey arrows indicate a significant mediation pathway. +/+: control, +/-: heterozygous *Drd4*. Asterisks indicate post hoc comparisons. \**p* < 0.05, \*\**p* < 0.01, \*\*\**p* < 0.001.

#### Females

A similar effect of condition was observed for puberty onset in females (F(2, 38) = 13.09, p = .003, Fig. 3f), where vaginal opening of LN reared mice occurred at a later stage than in SN or CN reared animals, irrespective of genotype (F(2, 38) = 0.28, p = .947). In contrast to males, female body weight at puberty onset was affected by rearing condition (F(2, 38) = 4.29, p = .010, Fig. 3g). Post-hoc testing revealed that CN reared females had increased body weight at the time of puberty onset. As in males, neither puberty onset (F(1, 39) = 0.52, p = .477) nor bodyweight at puberty onset (F(1, 39) = 0.09, p = .767) was affected by *Drd4* genotype. Similar to males, a negative correlation between body weight at weaning and puberty onset was observed (r = −0.46, p < .001, Fig. 3h). Mediation analysis revealed that body weight at weaning was a significant mediator of puberty onset for both LN (95%CI = [0.36, 1.66], Fig. 3j) and CN reared animals (95%CI = [−0.96, −0.08]). Although received entropy levels positively correlated with puberty onset in females (r = 0.31, p < .001, Fig. 3i), it did not mediate the effects of rearing condition on puberty onset (LN: 95%CI = [−1.21, 0.83]; CN: 95%CI = [−0.29, 0.23]).

### 3.4 Maternal care F1

No main effect of *Drd4* genotype was observed for any of home-cage maternal behaviors. However, mice that were exposed to different rearing conditions during early development displayed altered levels of arched-back nursing (ABN) towards their own offspring (F(2, 33) = 4.02, p = .027, Fig. 4a). LN reared dams performed less ABN than SN reared animals, irrespective of genotype (condition*genotype interaction: (F(2, 33) = 1.32, p = .275). Passive nursing levels were not affected by either condition (F(2, 33) = 0.73, p = .475, Fig. S2a) or the condition*genotype interaction (F(2, 33) = 0.58, p = .630). Rearing condition affected total nursing levels (F(2, 33) = 4.79, p = .024, Fig S2b), with an increase in CN reared mice compared to LN dams. Rearing condition also affected the time dams spent on the nest site (F(2, 33) = 7.40, p = .002, Fig. 2b), irrespective of genotype (F(2, 33) = 0.13, p = .845). Dams reared in the LN environment spent less time on the nest site compared to SN and CN reared animals. A main effect of rearing condition was also observed for the percentage of time dams spent licking/grooming their own pups (F(2, 33) = 4.51, p = .011, Fig. 4c), a key maternal behavior: LN reared dams spent less time licking/grooming than dams reared in a communal nesting environment. Moreover, while *Drd4* genotype did not affect LG levels (F(1, 34) = 0.10, p = .758), an interaction between genotype and rearing condition was observed (F(2, 33) = 4.99, p = .028). *Drd4*^*+/-*^ dams reared in the LN environment exhibited the lowest LG levels, whereas CN reared *Drd4*^*+/-*^ mice spent the most time licking/grooming their own pups.

**Figure 4.**
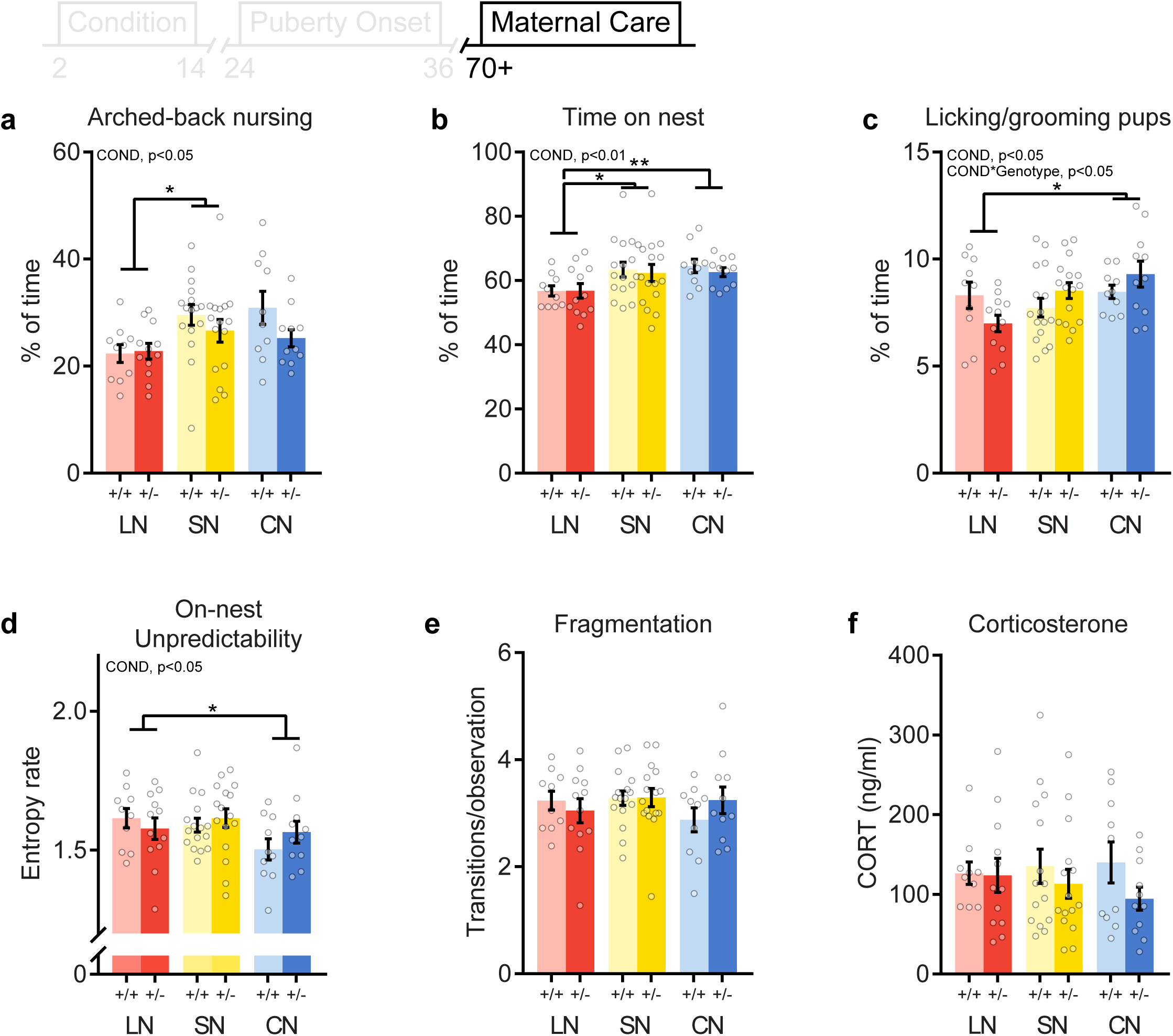
Effects of different rearing conditions and *Drd4* genotype on maternal care and basal corticosterone levels in female F1 offspring. Overall (P2-9) levels of (**a**) Arched-back nursing, (**b**) time on nest and (**c**) licking grooming exhibited by F1 female dams. (**d**) On-nest unpredictability and (**e**) fragmentation (on/off nest transitions) of maternal behavior. (**f**) Basal corticosterone levels. +/+: control, +/-: heterozygous *Drd4*. Group size: LN +/+: n = 10, LN +/-: n = 12, SN +/+: n = 16, SN +/-: n = 16, CN +/+: n = 10, CN +/-: n = 11). Asterisks indicate post hoc comparisons. \**p* < 0.05, \*\**p* < 0.01.

While F0 dams did not differ in total entropy rate, rearing condition had a significant effect on the total entropy rate of maternal behavior of F1 dams (F(2, 33) = 3.20, p = .032, Fig. S2c). Unpredictability was decreased in CN reared mice compared to dams reared in a SN environment. No genotype (F(1, 34) = 0.19, p = .733) nor interaction (F(2, 33) = 0.72, p = .411) effect was observed. In addition to the effects on total unpredictability, on-nest unpredictability was also affected by condition (F(2, 33) = 3.62, p = .044), where CN reared dams displayed lower rates compared to LN reared animals (Fig. S4d). On-nest unpredictability was unaffected by genotype (F(1, 34) = 0.31, p = .579) or the condition*genotype interaction (F(2, 33) = 0.85, p = .374). Fragmentation of maternal care was not affected by early life condition (F(2, 33) = 1.08, p = .269, Fig. 4e), genotype (F(1, 34) = 0.12, p = .728), or the interaction (F(2, 33) = 0.54, p = .505). Thus, while CN animals were raised with more fragmented maternal care, they did not differ in this behavior themselves when allowed to breed in a standard nesting condition. Cox regression revealed that pup retrieval was unaffected by rearing condition (hazard ratio 95%CI = [0.72, 1.39], p = 0.986), but *Drd4* genotype affected completion rate of pup retrieval (hazard ratio 95%CI = [1.03, 2.85], p = 0.040). *Drd4*^*+/-*^ dams were more likely to retrieve all pups within 5 minutes than *Drd4*^+/+^ animals (Fig. S2d).

Rearing condition of the dam affected body weight of the next generation (F2) at P2 (F(2, 33) = 4.39, p = .012, Fig. S2d), which was decreased in offspring from a LN reared mother compared to offspring from CN reared dams. However, this was normalized at weaning at P21 (F(2, 31) = 2.38, p = .313, Fig. S2e). Genotype of the dam did not affect offspring body weight (P2: (F(1, 34) = 0.06, p = .816); P21 (F(1, 32) = 0.52, p = .475)), nor did it interact with rearing condition (P2: (F(2, 33) = 0.01, p = .988); P21 (F(2, 31) = 0.97, p = .400)).

### 3.5 Corticosterone

Basal levels of blood plasma corticosterone were not affected by rearing condition (F(2, 32) = 0.10, p = .904, Fig. 2f) nor genotype (F(1, 33) = 2.04, p = .163). Moreover, *Drd4* genotype did not interact with rearing condition to affect corticosterone levels (F(2, 32) = 0.55, p = .532).

## 3. Discussion

In the present study, we examined the causal contribution of *Drd4* in differential susceptibility with a randomized experiment in rodents, allowing strict control for both genetic variation functioning –using *Drd4*^*+/-*^ mice- and early-life environmental factors. We demonstrate how different environmental conditions affect maternal care of mouse dams and subsequent sexual maturation in offspring. Interestingly, different rearing conditions during early development alter maternal care of adult mice towards their own offspring. Mice reared in a limited nesting/bedding environment are poor mothers in terms of arched-back nursing levels and nest presence. In contrast, communal nesting during early development results in mice that display lower rates of unpredictability in their own maternal behavior. Differential susceptibility was observed only for licking/grooming levels of female offspring towards their litter, of which LN and CN reared *Drd4*^*+/-*^ mice exhibited the lowest and highest levels, respectively.

### 4.1 Modelling impoverished and enriched rearing environments

The pattern of F0 maternal care resulting from exposure to the LN condition was largely in line with earlier findings using this model (Davis et al., 2017; Knop et al., 2019; Molet et al., 2016; Rice et al., 2008). While different pup-directed maternal behaviors remained relatively unaltered, the unpredictability of maternal behavior, particularly on the nest site, increased. However, in contrast to other reports, but in line with previous findings from our own lab (Knop et al., 2019) fragmentation of maternal care was similar to control conditions. In addition, it is important to note that pups in the LN condition were deprived from normal levels of licking/grooming upon first exposure to this condition on P2, whereas LG levels were similar to the SN and CN conditions from P3-P8. Moreover, a different circadian pattern in nest occupancy indicates that, similar to earlier results (Knop et al., 2019), LN dams exhibited altered circadian rhythmicity in maternal care, stressing the point that multiple time-points across the day-night should be examined to better grasp the implications of the LN condition.

Mouse dams adapted their maternal care to the communal nesting condition by decreasing nursing levels and increasing feeding behavior. However, despite decreased nursing time per dam, offspring body weight was similar compared to SN reared animals. This could be explained in part by the observation that pups in the communal nesting condition have increased accessibility to at least one mouse dam, a hallmark of the early social enrichment provided by this model (Branchi and Cirulli, 2014). In addition, litters in the CN condition are of a larger litter size, likely requiring less energy per pup to regulate body temperature.

### 4.2 Sexual maturation

The delayed puberty onset observed in both male and female LN reared mice was mediated by a decrease in body weight gain at weaning. The importance of body weight and leptin in regulating puberty onset is well-known for both humans (Lee et al., 2007; Tomova et al., 2015) and rodents (Ahima et al., 1997). We therefore also measured body weight at puberty onset for the adolescent mice that were raised in different early life conditions. The minimal differences in body weight at puberty onset suggest that, irrespective of early life background and subsequent body weight at weaning, the majority of mice postpone the onset of puberty until a certain body weight is reached. Only female mice reared in a CN setting showed increased body weight at puberty onset, indicating that these animals might exhibit, in line with the acceleration hypothesis, a relative delay in puberty onset, irrespective of bodyweight. It should be noted that early-life adversity not only affects body weight, it also alters adipose tissue, plasma leptin and leptin mRNA levels (Yam et al., 2017). Therefore, the mediation of puberty onset following LN is more complex and should be studied in more detail than only examining body weight per se. Nevertheless, the lack of differences in body weight at puberty onset between LN and SN reared mice, in combination with the delayed puberty onset of female mice that experienced increased unpredictability during rearing are not in line with the acceleration hypothesis of life history earlier proposed in humans. This may point to species differences but could also signify the relevance of uncontrolled factors in humans (e.g. caloric intake) that are controlled for in the current design.

### 4.3 Rearing conditions affect later-life maternal care

Different rearing conditions have been shown to affect maternal care provided to the next generation in the LN (Roth et al., 2009) and CN (Curley et al., 2009) models. Although previous results from our lab showed no effects of either LN or CN from P2-9 on adult maternal behavior (Knop et al., 2019), the results presented here do support long-lasting effects of rearing condition on maternal care. This could be explained by the duration and timing of exposure to early-life rearing conditions (P2-P9 in previous study compared to P2-14 in the present study). Given the different trajectories in brain circuit development (Hensch, 2005; Rice et al., 2000), the effects of early-life adversity, and potentially also enrichment, strongly depend on the critical period during which it occurs (Peña et al., 2019). The importance of this critical or sensitive period is highlighted by a recent study showing that different windows of exposure to the LN paradigm alter susceptibility to social defeat stress during adulthood (Peña et al., 2017). By extending the exposure of pups to different rearing conditions the development of brain regions involved in the regulation of maternal care, like the MPOA and mPFC (Dulac et al., 2014), may have been targeted more profoundly.

Extensive research from Meaney and co-workers have identified the pivotal role of arched-back nursing and licking/grooming behavior in rodent development (Caldji et al., 1998; Liu et al., 1997; Meaney, 2001). Many studies investigating intergenerational transmission of maternal care observe a similar phenotype in the offspring and the mother (Champagne, 2008; Curley et al., 2008). Interestingly, the lower ABN and nest occupancy levels of LN reared female mice observed in our current study did not coincide with a lower ABN or nest presence of their own mother. On the contrary, female LN reared pups experienced *increased* levels of nest occupancy by the dam compared to the SN condition, but showed *lower* levels of nest occupancy when taking care of a litter themselves. Similarly, CN reared mice received comparable levels of unpredictability as standard reared mice, yet provided more predictable maternal behavior towards their own offspring. Finally, LN reared animals received increased on-nest unpredictability but showed similar on-nest entropy rates compared to SN reared dams. Thus, although the differences in maternal care of F1 dams presented here are not mimicking the phenotype of the mother, the quality of the early-life environment (poor vs. enriched) did affect the quality of F1 maternal care under standard breeding conditions.

For licking/grooming behavior, the effects of rearing conditions were restricted to *Drd4*^*+/-*^ animals, whereas rearing conditions had no effect on LG levels in wild-type animals. Using *Drd4* genotype as a susceptibility factor, this is supportive evidence for differential susceptibility in our controlled animal model with respect to a key feature of rodent maternal care, across generations. Studies on differential susceptibility in humans focused predominantly on the effects of maternal care on child development, highlighting the increased susceptibility of *DRD4-7R* carrying children to parental sensitivity (Bakermans-Kranenburg et al., 2008a). However, these studies have not yet examined parental care of the next generation.

Clearly, the exact mechanisms through which the early-life environment impacts on later-life behavior remain to be elucidated. Previous studies suggest an important role for the methylation of genes involved in the HPA-axis (Turecki and Meaney, 2016). Human studies also link the *DRD4-7R* genotype to alterations in components of the HPA-axis. Gene-early environment effects have been observed for basal cortisol in children (Bakermans-Kranenburg et al., 2008a), as well as stress induced cortisol levels of young adults (Buchmann et al., 2014). A prominent role for alterations in circulating basal corticosterone levels in adulthood is not supported by our data. However, stress reactivity was not assessed and could, at least in part, underlie the observed alterations in maternal care.

Other systems may also be critical in the mechanism underlying differential susceptibility. Recent studies using different molecular tools and mouse knock-in models have begun to unravel the exact function of the *DRD4-7R* in corticostriatal glutamatergic neurotransmission, enhancing our understanding of the *DRD4* receptor and susceptibility to the environment (Bonaventura et al., 2017; González et al., 2012). Other studies used a wide array of novel techniques to show the involvement of dopamine receptors in mediating the social deficits observed after severe early-life stress (Shin et al., 2018). These advances in our understanding of the functioning of different dopamine receptors in regulating susceptibility will help to guide future studies into the role of *DRD4*.

There is increasing awareness that most consequences of early-life rodent models have small effect sizes (Bonapersona et al., 2019), which is also the case in our study. Although we have very decent group numbers compared to common practice in this field, we should take this into consideration and interpret the results with care. To increase statistical power in future experiments, animal numbers should be adapted to realistically expected effect sizes and animal ethical committees should be aware of this (Button et al., 2013). Moreover, meta-analyses in this field should be stimulated and can help in designing future studies (Bonapersona et al., 2019).

## Conclusion

The research presented here provides a translational approach to examine the contribution of the *Drd4* gene in differential susceptibility. While other preclinical studies on differential susceptibility in socially monogamous prairie voles focused on the role of prenatal stress in enhancing developmental plasticity to both adverse and supportive contexts (Hartman et al., 2018; Hartman and Belsky, 2018), we show that adverse or enriched postnatal environments also interact with genetic factors in mice, for better and for worse. Future experiments should be targeted to test which neurobiological mechanisms are involved in mediating the effects of *DRD4* with regard to differential susceptibility.

## Acknowledgements

This work was supported by the Consortium on Individual Development (CID), which is funded through the Gravitation program of the Dutch Ministry of Education, Culture, and Science and the Netherlands Organization for Scientific Research (NWO grant number 024.001.003). M.J. Bakermans-Kranenburg was supported by the European Research Council (ERC AdG 669249); M.H. van IJzendoorn was supported by the Netherlands Organization for Scientific Research (Spinoza Prize).

A preprint version of this article was uploaded at bioRxiv (www.biorxiv.org)

---

**Figure S1.**
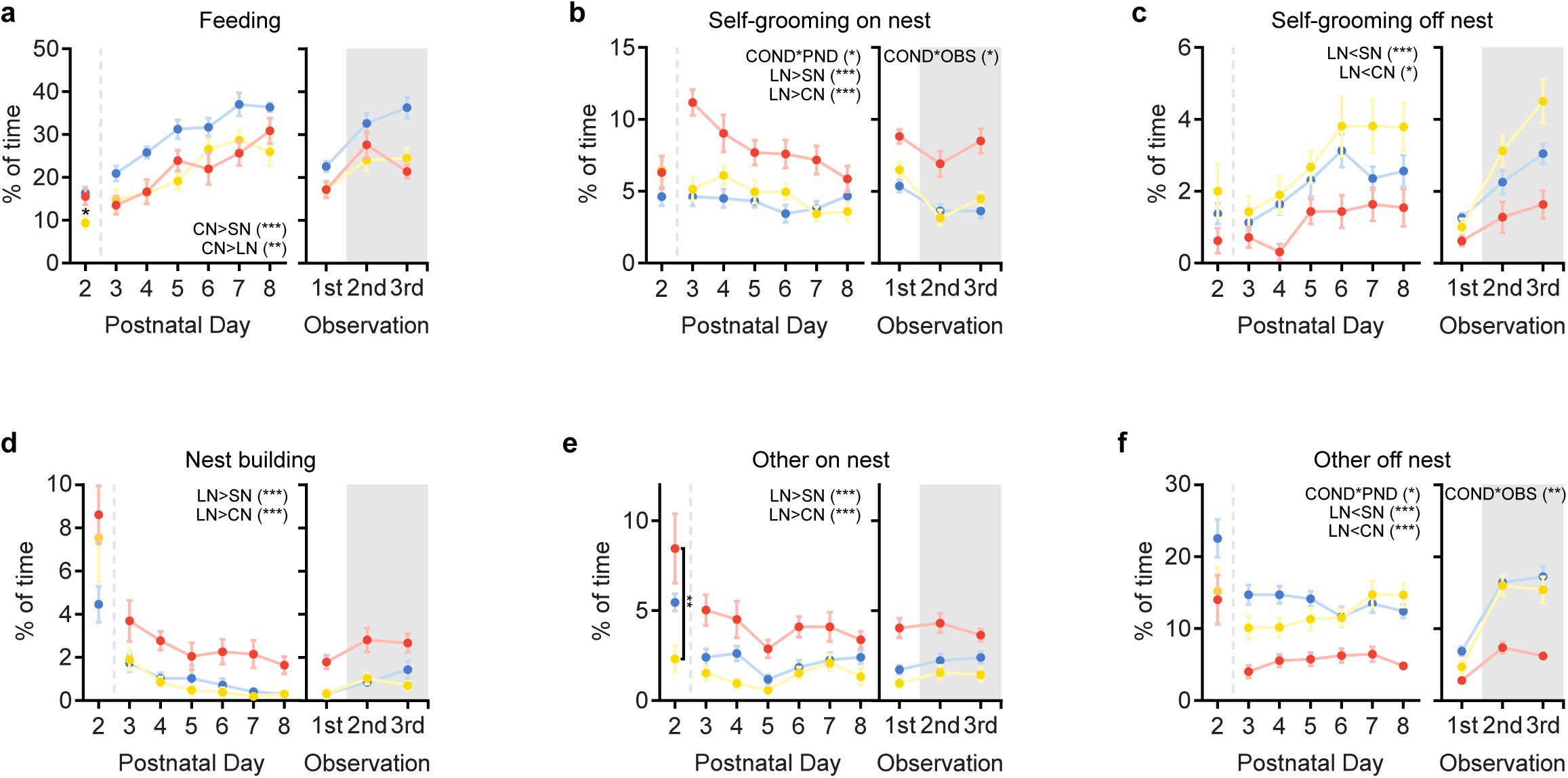
Effect of different housing conditions on maternal care. (**a**) Feeding, (**b**) self-grooming on nest, (**c**) self-grooming off nest, (**d**) nest building, (**e**) other on nest and (**f**) other off nest behavior for limited nesting (red, n = 13), standard nesting (yellow, n = 14) and communal nesting (blue, n = 13) dams, depicted over postnatal days (left) and time of the day (right). The shaded area indicates the dark phase of the LD cycle. Statistics indicate main effects or interactions. \**p* < 0.05, \*\**p* < 0.01, \*\*\**p* < 0.001.

**Figure S2.**
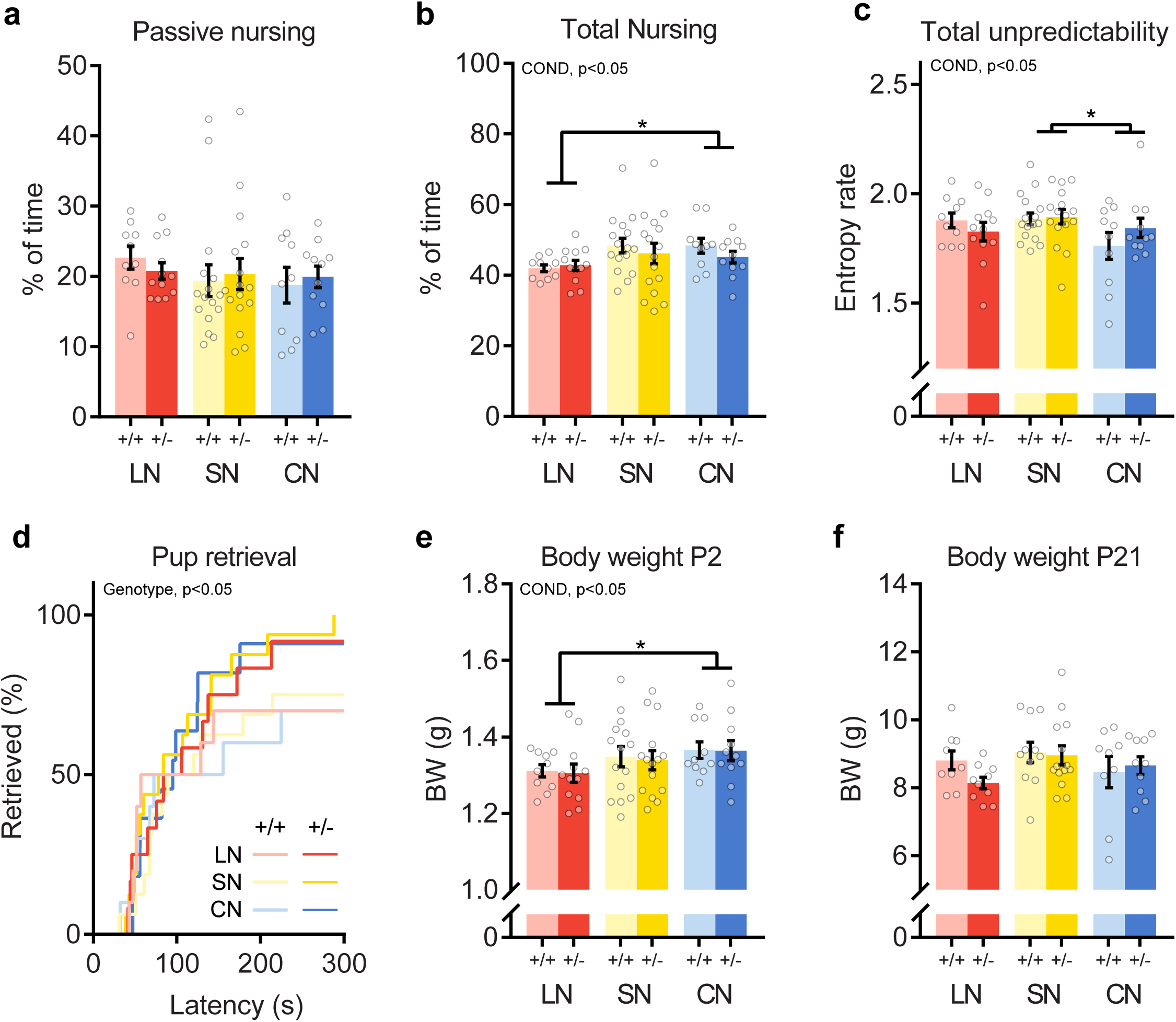
Effects of different rearing conditions and *Drd4* genotype on F1 maternal care and F2 body weight. (**a**) Passive nursing and (**b**) total nursing levels. (**c**) Total unpredictability rates. (**d**) Pup retrieval latencies and completion rates. (**e**) F2 offspring body weight at P2 and (**f**) P21. +/+: control, +/-: heterozygous *Drd4*. Group size: LN +/+: n = 10, LN +/-: n = 12, SN +/+: n = 16, SN +/-: n = 16, CN +/+: n = 10, CN +/-: n = 11). Asterisks indicate post hoc comparisons. \**p* < 0.05, \*\**p* < 0.01.

